# Spatial structure undermines parasite suppression by gene drive cargo

**DOI:** 10.1101/728006

**Authors:** James J Bull, Christopher H Remien, Richard Gomulkiewicz, Stephen M Krone

## Abstract

Gene drives may be used in two ways to curtail vectored diseases. Both involve engineering the drive to spread in the vector population. One approach uses the drive to directly depress vector numbers, possibly to extinction. The other approach leaves intact the vector population but suppresses the disease agent during its interaction with the vector. This second application may use a drive engineered to carry a genetic cargo that blocks the disease agent. An advantage of the second application is that it is far less likely to select vector resistance to block the drive, but the disease agent may instead evolve resistance to the inhibitory cargo. However, some gene drives are expected to spread so fast and attain such high coverage in the vector population that, if the disease agent can evolve resistance only gradually, disease eradication may be feasible. Here we use simple models to show that spatial structure in the vector population can greatly facilitate persistence and evolution of resistance by the disease agent. We suggest simple approaches to avoid some types of spatial structure, but others may be intrinsic to the populations being challenged and difficult to overcome.

## INTRODUCTION

Genetic engineering has advanced to the point that it is not only possible to introduce arbitrary, massive changes into the genomes of countless organisms, but it is also possible to engineer changes that rapidly sweep throughout an entire species. The rapid sweeps are enabled by a class of genetic elements called gene drives that function on the principle of biasing transmission in gametes or in survival (Hamilton, 1967; Lyttle, 1977; Burt, 2003; Deredec et al., 2008; Gould, 2008; Gould et al., 2008; Report, 2016). Perhaps the most powerful use of a gene drive is one that suppresses population numbers and even potentially drives the population extinct. A more benign form of gene drive is one that sweeps without causing much harm to its host. Any gene drive may be harnessed with additional genetic material (i.e., ‘effector gene’ or simply ‘cargo’) that is carried along with the drive as it spreads (Sandler and Novitski, 1957; Gould, 2008; Gould et al., 2008; Gantz et al., 2015). A harmless drive equipped with a cargo provides a fast and simple means of genetically transforming a population, potentially endowing that population with properties that meet social goals without harming the species.

The application of gene drives is limited in a few important ways. They require species with largely obligate sexual reproduction and moderate to high rates of outcrossing. Furthermore, drives that impair fitness are highly susceptible to evolution of resistance (Burt, 2003; Noble et al., 2017; Unckless et al., 2017; Bull et al., 2019). For these reasons, some applications are most amenable to species modification with harmless gene drives carrying genetic cargo. One such application is the use of gene drives to transform disease vectors so that the disease agent (‘parasite’ or ‘pathogen’) can no longer be transmitted: the pathogen cannot be targeted with a gene drive, but its vector can. Such a gene drive can be designed to have little effect on the vector yet completely block the pathogen (Sandler and Novitski, 1957; Burt, 2003; Gantz et al., 2015).

Multiple approaches to population replacement that involve gene drive have been proposed. Several of these have been implemented successfully in model systems, and important progress has been made with homing-based approaches. Current technology using CRISPR homing drives appears good enough to allow a gene drive to avoid resistance evolution and achieve wide coverage of a population (Kyrou et al., 2018; Champer et al., 2019c). Thus, an inhibitory cargo should also be able to achieve wide population coverage and thereby eradicate many types of parasite, provided that the cargo’s suppression of the parasite cannot be overcome by single mutations. One potential limitation of this approach is that even slight fitness costs of cargo carriage will ultimately lead to a decay of cargo in the vector population, but the decay should often be slow enough to allow parasite suppression for tens to hundreds of generations (Beaghton et al., 2017) – still potentially enough for eradication.

Here we suggest another possible basis of cargo failure, spatial structure in the host population combined with imperfect gene drive coverage/expression. If parasite movement is limited, even small areas of incomplete suppression may allow parasite persistence that serve as nuclei for evolution of parasite resistance. We offer simple models of the sensitivity of parasite persistence and resistance evolution under spatial structure to gauge the plausibility of parasite escape from gene drive control. Our approach potentially applies to any widespread genetic modification of a population, not just gene drives.

## BACKGROUND

This section offers a biological framework for the problem addressed in the models section. This framework is easily explained at an intuitive level and helps anticipate the models. We henceforth use ‘parasite’ instead of ‘pathogen,’ to avoid confusion as to the effect of the parasite—it will commonly be a pathogen of humans but not necessarily of the vector, where it is targeted.

### Two kinds of engineered gene drives

The gene drives proposed and developed for genetic engineering fall into two classes. One class relies on homing, whereby the drive element cuts the genome at a specific site and inserts itself into that site (Burt, 2003; Gantz and Bier, 2015; Kyrou et al., 2018). A homing drive’s fitness advantage comes from a transmission bias in gametes of heterozygotes. CRISPR technology has greatly facilitated this type of engineering because CRISPR-Cas9 is a site-directed nuclease. The other class relies on biased offspring survival, many of which are known as ‘killer-rescue’ systems (Chen et al., 2007; Gould et al., 2008; Marshall and Hay, 2011; Legros et al., 2013; Akbari et al., 2013, 2014; Buchman et al., 2018; Oberhofer et al., 2019; Champer et al., 2019a). One of the major differences between these two classes of drive elements is the speed and ease with which they spread. A homing element spreads rapidly and can, in theory, be successfully introduced with a single individual. Killer-rescue systems spread more slowly and often must be introduced above a threshold density to spread, although that distinction is not absolute (Champer et al., 2019a).

### Mass action dynamics

Gene drives have traditionally been modeled and understood in the context of well-mixed populations (e.g., Prout, 1953; Bruck, 1957; Hamilton, 1967; Burt, 2003; Marshall and Hay, 2011; Legros et al., 2013; Akbari et al., 2014; Unckless et al., 2015; Beaghton et al., 2017; Godfray et al., 2017). A homing gene drive gains its advantage from heterozygotes, the non-drive allele of a heterozygote being replaced with the drive allele during reproduction. Heterozygote frequency is enhanced with outcrossing (mass action), depressed with inbreeding and some other types of assortative mating. Even killer-rescue systems presumably rely on mixing, so that the killer and rescue alleles are not closely associated, and the rescue allele thereby gains a large benefit. With mass action, an efficient homing drive can spread from low frequencies to near-fixation in close to 10 generations (Burt, 2003; Godfray et al., 2017; Beaghton et al., 2017).

The evolution of a gene drive and its associated cargo can be divided into two phases. The first phase encompasses the short-lived spread of the drive. Although gene drives are potentially highly efficient, various types of fitness effects, imperfections in the drive mechanics and variation in the host population can limit the final coverage of the drive (Deredec et al., 2008; Godfray et al., 2017; Beaghton et al., 2017; Champer et al., 2017). Once the drive has spread to its limit, phase two sets in, whereby evolution proceeds according to fitness effects on the host. Any fitness cost stemming from the drive allele or its genetic cargo now begins to select a population reversal toward loss of the cargo and/or drive, favouring alleles resistant to the drive or cargo-free drive states. The speed of this reversal depends heavily on fitness costs and on the initial frequencies of the different parties; it is typically much slower than the spread of the drive (Beaghton et al., 2017). In the long term, a genetic cargo with any fitness cost will be lost. The social benefits of the cargo must therefore be manifest in a time frame compatible with its expected duration.

### Population spatial structure

Gene drives require reproduction. Their spread will thus follow the conduits of reproductive connections in the host population, which may well have a strong spatial component—as when individuals mate with neighbors (North et al., 2013; Beaghton et al., 2016; Tanaka et al., 2017). Any genetic variation that arises in the gene drive or cargo, such as mutations that delete or down-regulate the cargo, will be propagated along those conduits and expand accordingly, leading to spatial structure in parasite inhibition (Beaghton et al., 2017). Even more simply, for purely dynamical reasons, the drive may fail to reach isolated pockets of the population (North et al., 2013). In turn, spatial structure of a genetically variable inhibitor will often mean that different locations of the parasite experience different levels of inhibition. With spatial structure, even small regions of reduced parasite suppression may enable parasite persistence which then facilitate parasite evolution of resistance to the inhibitor.

Pre-existing genetic variation in the host population may also affect gene drive efficacy, spread and cargo expression (e.g., Drury et al., 2017; Champer et al., 2019b). For example, some designs for a harmless homing drive have it target a genomic region that can be disrupted with little or no fitness effects; such a region may thus not be strongly selected to conserve sequences and may be variable across the host population, blocking gene drive spread in some regions. (One design avoids this problem by targeting an essential gene and carrying a cargo that replaces the targeted gene (Burt, 2003; Champer et al., 2019c).) Cargo gene expression may likewise be affected by the genome in which it resides, and geographic variation in genomic content may lead to geographic variation in cargo expression.

Our intent is to investigate the consequences of spatially structured inhibition of the parasite/pathogen. The details of structure will typically be implementation-specific, but an appreciation for the importance of spatial structure when it exists may be a requisite for successful application of a gene drive cargo.

## ANALYSIS

### Mathematical models

Our model is most easily applied to an asexual pest/parasite infecting a single host species. Although not specifically modeled, our problem may be extended in spirit to a parasite transmitted between two host species, as to *Plasmodium* transmitted by a mosquito to humans and back to mosquitoes; in this case, the gene drive is introduced into the mosquito to block *Plasmodium* reproduction and transmission. Our models merely omit the second host, but we conjecture that the effects they reveal apply to that case, subject to some conditions mentioned in the Discussion.

The social goal is to suppress parasite reproduction with a genetic cargo in the host. To keep the problem simple, we assume that a gene drive and its cargo have already swept through the host species. (The actual process of gene drive evolution is thus ignored, and indeed, non-gene-drive methods of cargo infusion may also be used to achieve this end (Okamoto et al., 2014).) In any one host individual, parasite inhibition by the cargo occurs in one of three states: (i) full inhibition, (ii) partial inhibition, or (iii) no inhibition. Partial inhibition would result from weak expression of the cargo in the host; no inhibition would result from loss of the cargo from the gene drive or resistance to the gene drive itself, such that the host individual lacks the gene drive and its cargo altogether. The formulation of the model is trivially extended to multiple states of partial inhibition.

The model counts numbers of parasites in each type of host. Host type merely translates into parasite reproduction. Notation is

*x*_*h*_: The relative frequency of hosts of type *h*

*b*_*gh*_: The fecundity of a parasite genotype *g* in hosts of type *h*

*n*_*gh*_: The current number of parasites of genotype *g* in hosts of type *h*

*N*_*g*_: The current number of progeny produced by genotype *g* across all patches

*α*: The fidelity of parasite reproduction to hosts of the same type.

where parasite genotype *g* ∈ {0, 1, *…*, *M*} and host type *h* ∈ {0, 1, *…*, *H*.} Here, host type 0 indicates hosts with no cargo (hence no parasite suppression), and larger values of *h* correspond to increased levels of suppression, with *H* denoting the number of types of cargo-carrying hosts differing in some aspect of parasite suppression; parasite genotype 0 is the wild type (with no protection against the cargo), and larger values of *g* correspond to mutant strains with increased levels of resistance to the cargo.

To approximate the separation of phases between rapid gene drive spread and the subsequent effect of cargo on the parasite, we let the *x*_*h*_ and *b*_*gh*_ be constant in time, so the only changes are in parasite numbers. Time is discrete. For biological reasons, in the case of three host types, we also assume that the fecundity of parasite genotype 0 satisfies *b*_00_ > 1 > *b*_01_ > *b*_02_, which means that the parasite has negative growth in all host types except the one lacking cargo.

To set the stage for a structured host population, we suppose that hosts are clustered in *patches* of similar host types, patch types designated by subscript *h*. (’Host’ and ‘patch’ are used interchangeably below, but ‘patch’ helps convey structure.) Patches could result, for example, from limited host migration and gene drive spread through the host population in a manner that follows host population structure. The clustering of hosts and the consequent movement of parasites between patches determines the extent to which structure is experienced by the parasites. To establish a mass-action baseline, adult parasites reproduce and release all progeny into a random pool, from which they settle into each of the *H* + 1 patch types at frequencies *x*_0_, *x*_1_, *…*, *x*_*H*_.

Spatial structure is modeled indirectly by assuming that a fraction *α* of the progeny born in a patch type remains in the same patch type without entering the random pool; this ‘fidelity’ increases with the retention of progeny in their natal patch type. This process is fundamentally the same as migration in standard population genetics problems (Crow and Kimura, 1970). Our formulation is different in that *α* denotes a lack of movement from the natal site instead of movement between patches/populations; this formulation leads naturally to calculating the null case of mass action (*α* = 0), which is presumably the default expectation in gene drive applications. Note that fidelity to a patch type is imposed without inbreeding, so the numbers within a patch are assumed large enough that consanguinity can be ignored.

We first consider a simple system of linear dynamical equations. The overall progeny output of strain *g* across all environments is

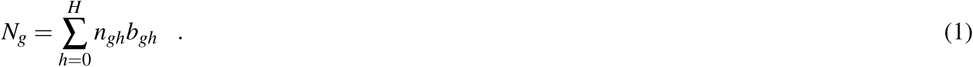

Using primes to indicate one generation hence,

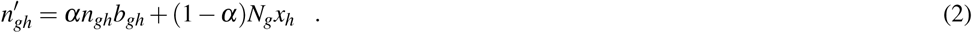

The joint dynamics for genotype *g* across all host types can be written as a matrix projection recursion (Caswell, 2006). Dropping the genotype subscript *g* for ease of visualization, the recursion has form

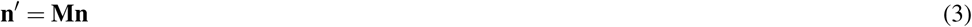

where **n** and **n**′ are (*H* + 1)-dimensional column vectors with *h*th element *n*_*h*_ and 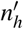, respectively. The (*H* + 1) × (*H* + 1) matrix **M** has diagonal elements *M*_*hh*_ = *b*_*h*_[*α* + (1 − *α*)*x*_*h*_] and off-diagonal elements *M*_*hi*_ = *b*_*i*_(1 − *α*)*x*_*h*_, *i* ≠ *h*. In words, matrix element *M*_*hi*_ describes the rate that individuals who originate in patch type *i* contribute to the abundance in patch type *h* at the next time step. The densities of the genotype *g* at any time *t*, **n**(*t*), can be computed as **n**(*t*) = **M**^*t*^ **n**(0) (Caswell, 2006).

This model allows for a biological anomaly: when a genotype *i* is initially assigned to a patch (*j*), a sufficiently large combination of fecundity and fidelity (*b*_*ij*_*α* > 1) allows its numbers to persist even when the patch type is absent (even when *x*_*j*_ = 0). This effect occurs because, once a genotype exists in a patch (by initial conditions), a portion of its growth comes from offspring who stay in the patch to reproduce. This effect is independent of patch size. Because we interpret *x*_*j*_ as the fraction of hosts of type *j*, we require *x*_*j*_ > 0 for any host type that harbors parasites. This requirement is further imposed when enforcing a carrying capacity (see below).

These equations assume fixed fecundities and thus allow unlimited population growth. They should be adequate to decide whether parasites persist or die out, because they can be applied to deterministic dynamics at low population densities to describe the direction of population growth, when fecundities are intrinsic and unaffected by densities. For dynamics and evolution in high-density populations, we use a related model that imposes density regulation. For simplicity, we limit our description of density regulation to the case of two patches and two strains.

### Introducing density regulation

Density regulation may be important in the evolution of alternative genotypes in a persisting population, at least because a small portion of the environment with cargo-free hosts may be limited in how many parasites it can support – a small patch may allow parasite persistence but have little impact on parasite numbers across the entire environment. We thus introduce a simple form of patch-specific density regulation that will be used in some numerical trials of two patch types, 0 and 1.

Let *K* be the overall upper density limit for the environment and let *K*_0_ = *Kx*_0_, *K*_1_ = *Kx*_1_ denote the ceilings for patch 0 and 1, respectively. Let Π_1_ = min *{n*_01_*b*_01_ + *n*_11_*b*_11_, *K*_1_ *}* be total parasite progeny production emanating from patch 1, and Π_0_ = min *{n*_00_*b*_00_ + *n*_10_*b*_10_, *K*_0_ *}* be the total parasite progeny production from patch 0, each limited locally, without respect to regulation in the other patch type. The extension to more than two patch types is straightforward.

To decide how overall parasite production in a patch is divided between two genotypes when progeny output is limited by carrying capacity, let *p*_11_ = *n*_11_*b*_11_*/*(*n*_01_*b*_01_ + *n*_11_*b*_11_) denote the fraction of offspring that are strain 1 within patch 1, and *p*_01_ = 1 − *p*_11_ the fraction in patch 1 that are strain 0. Similarly, in patch 0 we have fractions *p*_10_ = *n*_10_*b*_10_*/*(*n*_00_*b*_00_ + *n*_10_*b*_10_) and *p*_00_ = 1 − *p*_10_. Accounting for mixing and assuming both genotypes are density-regulated the same, the total number of genotype 1 entering patch 1 is

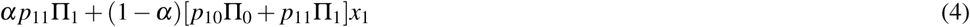

and the total number of genotype 0 entering patch 1 is

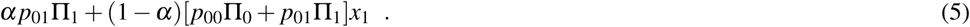

Analogous equations apply to genotypes 0 and 1 entering patch 0. Note that mixing can lead to a temporary violation of local carrying capacities when patches are repopulated by adults, but the capacity limit is imposed again at the next round of reproduction.

### Parasite persistence is facilitated by spatial structure

The growth or suppression of the parasite depends on whether the magnitude of the leading eigenvalue of **M** in eqn (3) exceeds 1. Density dependence can be ignored when addressing persistence, but an eigenvalue exceeding 1 merely indicates that parasites have positive growth somewhere in the environment, perhaps in only a tiny locale, with negative growth everywhere else. Although the characteristic equation is easily found, the leading eigenvalue (*λ*_*max*_) of a genotype is tractable for arbitrary *H* only for the extremes of mass action (*α* = 0), and complete parasite isolation among host types (*α* = 1). For mass action,

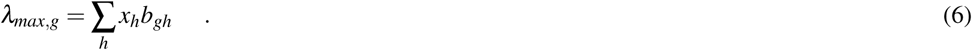

As is well appreciated for mass action, parasite growth is just the weighted average fecundity across all host types. Small levels of weak suppression (i.e., low values of *x*_0_ despite possibly high values of *b*_00_) will not themselves enable parasite persistence except when parasite fecundity is extraordinarily high. When more than one genotype has an eigenvalue greater than 1, density dependence will determine which one prevails (see below).

With complete parasite separation across the host types (*α* = 1),

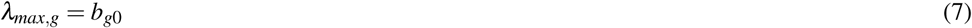

(*λ*_*max*_ is associated with patch type 0 because cargo-free hosts are assumed to offer the highest fecundity of all patches, regardless of genotype). Here, the parasites inhabiting each host type have their own eigenvalue, and any host type with *b*_*gh*_ > 1 will allow parasites of genotype *g* to persist in that patch type (subject to competition among different parasite genotypes). In this extreme, the values of *x*_*h*_ no longer matter: even a small fraction of permissive hosts will allow the parasite to persist.

The question motivating our study is how sensitive parasite persistence is to fidelity *α*—reflecting spatial structure. As the eigenvalue for several patch/host types is unwieldy, we reduce the problem to just two patch types, *g* = 0 and 1 (fully permissive and fully blocking of wild-type); this reduction also simplifies patch type abundances: *x*_0_ + *x*_1_ = 1. For this case, the largest eigenvalue is

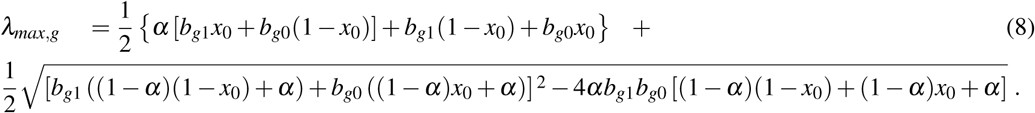

Values of the eigenvalues for different fecundities in the two patch types are shown in Fig. 1, the panels differing in fidelity (*α*) and cargo-free patch size (*x*_0_) values. The graphs show four contour lines, but *λ*_*max*_ = 1—the boundary between persistence and extinction—is thicker than the others. It is clear that persistence is enhanced by spatial structure, though typically a large cargo-free fecundity is required if the cargo is effective (*b*_00_ must be well above 1 when *α* and *b*_01_ are small). But for small cargo-free patch sizes (small *x*_0_), parasite persistence becomes possible despite low fecundity values in cargo-free hosts when the cargo becomes moderately ineffective (when *b*_01_ exceeds 1). In addition to the effect of fecundity, there are also effects of *α* and *x*_0_; one interesting effect is that the isoclines are visually step-like except in the lower left panel. For those cases, fecundity in cargo-bearing hosts has little effect on the parasite growth rate until *b*_01_ *≈ λ*_max_.

**Figure 1.**
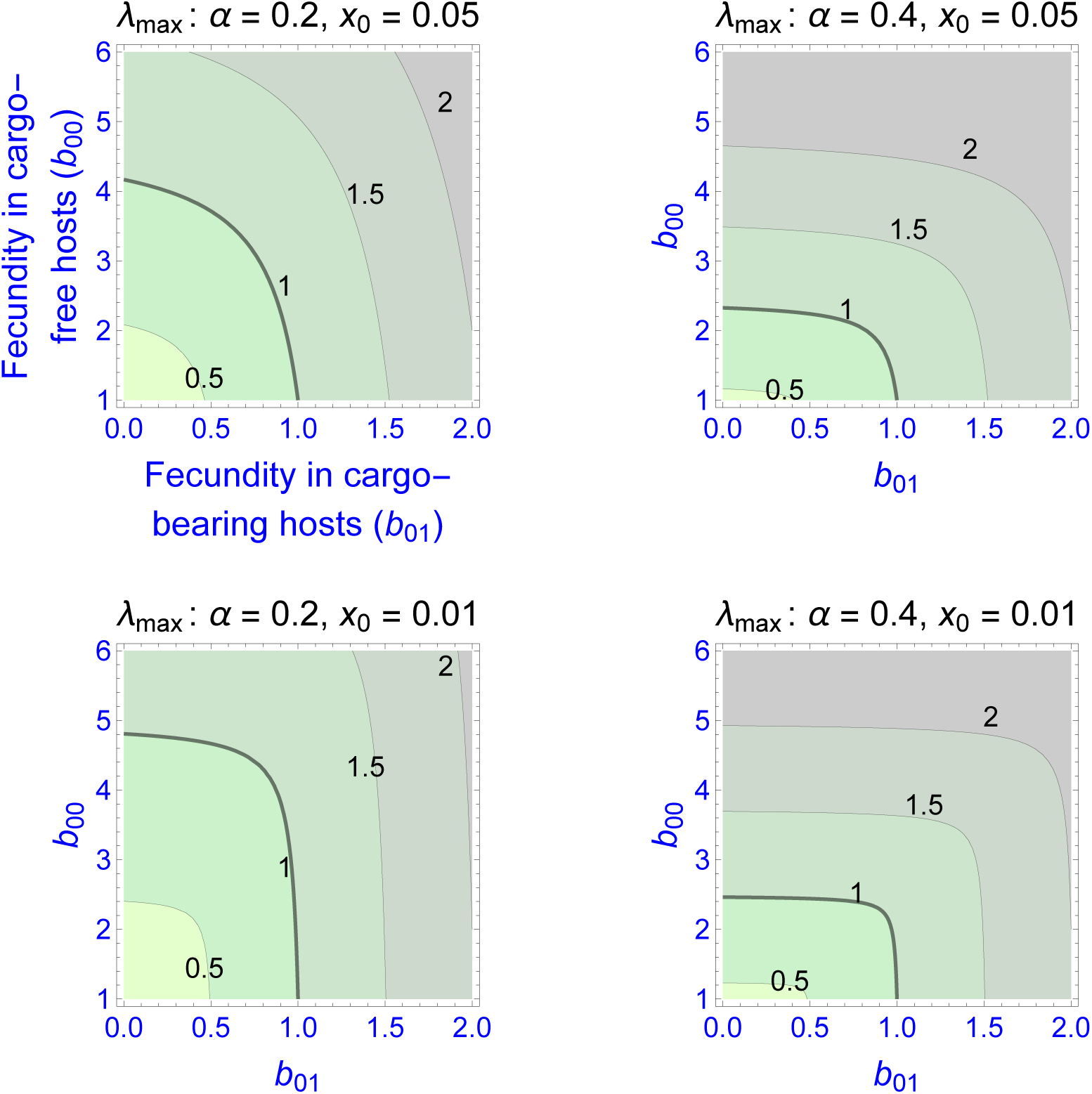
Contour plots of wild-type parasite growth rates, as given by eigenvalues (*λ*_*max*_, from eqn (8)). Each panel varies parasite fecundities in the two patch types (*b*_00_ is cargo-free fecundity, *b*_01_ is fecundity in cargo-bearing hosts. Parasite growth rates rise with increases in each fecundity, but the eigenvalues often show a step-like pattern in which fecundity increases in one host type have little effect until it reaches a threshold. Panels differ in parasite fidelity (*α*) and size of the cargo-free patches (*x*_0_); *λ*_*max*_ values are given adjacent to the contours. The wild-type genotype is represented as *g* = 0. Colors merely distinguish the regions bounded by the curves.

Most empirical interest is likely to be in the extreme case that the cargo completely suppresses wild-type parasite reproduction (*b*_01_ = 0), as that would be the goal of the engineer. Indeed, any gene drive release could be avoided until such an appropriate inhibitor was found. This case corresponds to the the sliver defined by the vertical axes in Fig. 1. Furthermore, this case is highly tractable:

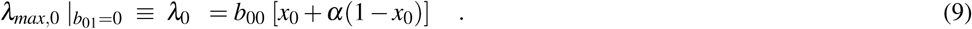

Persistence in this case requires *λ*_0_ = *b*_00_ [*x*_0_ + *α*(1− *x*_0_)] ≥ 1. This implies, for example, that persistence of the parasite is assured even when completely inhibited by the cargo as long as fecundity in cargo-free hosts (*b*_00_), exceeds [*x*_0_ + *α*(1 − *x*_0_)]^*-*1^. Consideration of this minimum cargo-free fecundity (Fig. 2) shows that spatial structuring with fidelity (*α*) well below 1 (e.g., 0.5) enables parasite persistence for even rare cargo-free hosts (small *x*_0_), as long as the parasite can grow there moderately well (e.g., *b*_00_ > 2). The results are largely insensitive to patch size *x*_0_ when fidelity reaches 0.6.

**Figure 2.**
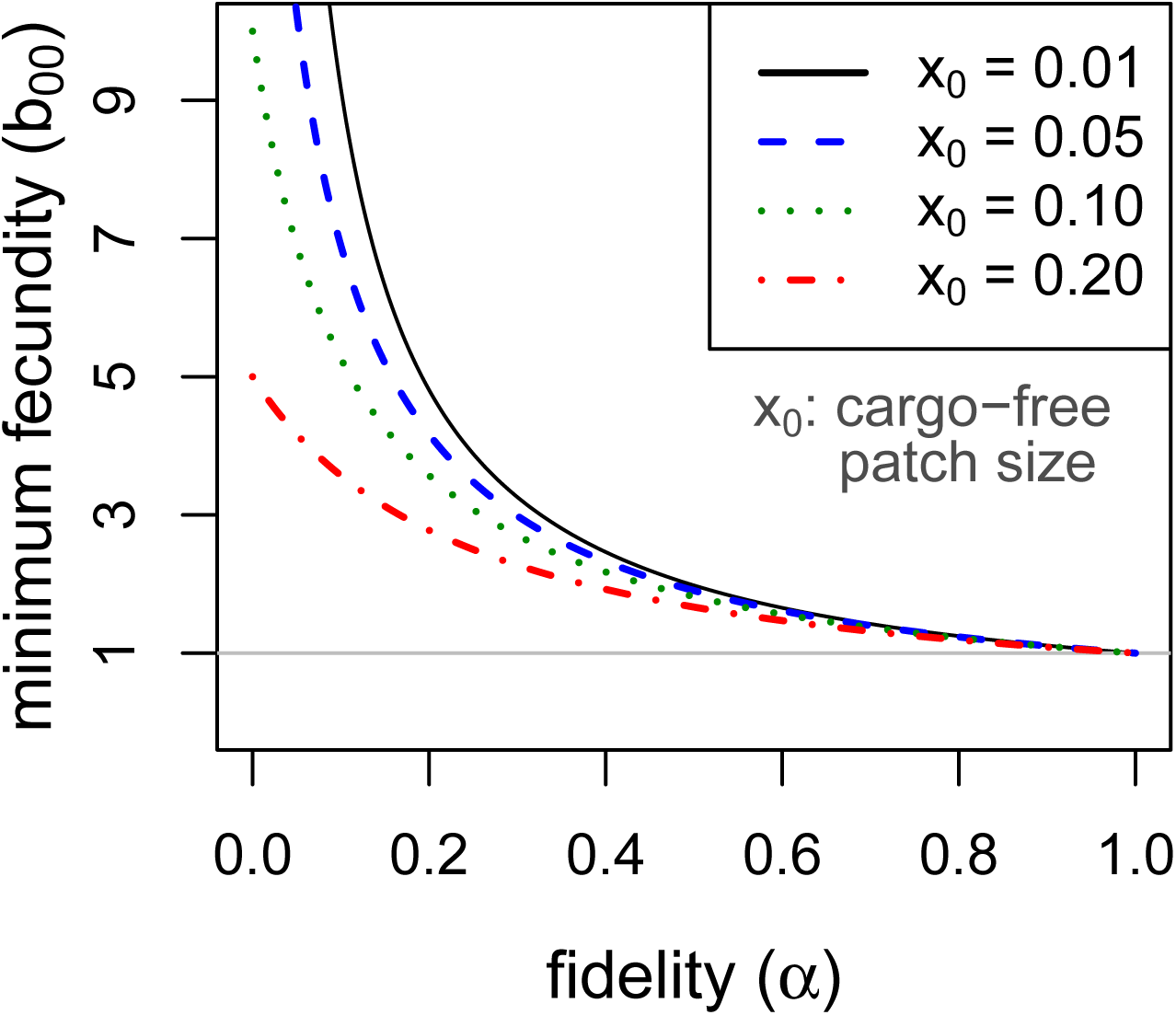
Minimum fecundity in cargo-free hosts required for parasite persistence when the cargo causes complete inhibition, (*b*_01_ = 0, from eqn (9)). Each curve represents a different size of cargo-free patch (*x*_0_); the required cargo-free fecundity for parasite persistence (*b*_00_, vertical axis) decreases with the fidelity to patch type (*α*, horizontal axis). For *α* ≥ 0.6, there is little effect of patch size. The curves intersect *α* = 0 at 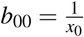. The horizontal line at *b*_00_ = 1 indicates the minimum fecundity required for the parasite to persist in the absence of cargo, which all curves intersect at *α* = 1.

It is also easy to see that, when cargo-free patch size *x*_0_ is small, as it should be if the engineering worked as expected, the effect of increasing fidelity *α* is approximately the same as increasing the frequency of permissive hosts (*x*_0_) – both are mostly linear effects and of the same magnitude. Whereas *x*_0_ is somewhat under the control of the engineer, *α* is not and has the potential to thwart parasite eradication. Thus, a very small *x*_0_ could easily cause parasite population collapse in the mass action case, but *b*_00_*α* > 1 is sufficient for persistence no matter how small *x*_0_. Persistence could be achieved by a cargo-free patch merely large enough that parasite progeny often did not disperse beyond the patch edges.

### Ease of resistance evolution depends on ecology

At the population level, the evolution of resistance to a genetic cargo has many parallels with the evolution of resistance to antibiotics and pesticides. The latter problems are thoroughly studied, and it is well appreciated that resistance is especially prone to evolve under intermediate levels of drug/chemical application (e.g., Gould and MacKenzie, 2002; Andersson and Hughes, 2012; Tabashnik and Gould, 2012; Neve et al., 2014; Gould et al., 2018). Inhibition by a gene drive cargo is different from chemicals in that the levels of inhibition are established at fixed, semi-permanent levels in the near term. They are also largely unchangeable, at least in the short term, should it be discovered that they are inadequate.

Of the many factors to consider, an important one is the mutational spectrum of resistance: a cargo for which simple, single mutations can allow parasite persistence seems doomed to fail, and intuition suffices for preliminary understanding, at least deterministically. Our interest instead lies in gradual evolution and the selection of weak resistance mutations. It might be hoped that cargo-based inhibitors can be found for which resistance mutations are impossible, but a more realistic hope is that inhibitors could be found for which resistance can evolve only gradually.

The evolution of resistance can be considered in two contexts. One is known as ‘evolutionary rescue,’ whereby the population is in decline and a resistance mutation potentially reverses the decline (Martin et al., 2013; Uecker et al., 2014; Hufbauer et al., 2015; Gomulkiewicz et al., 2017). The other context, the one addressed here, is resistance evolution in a persisting population. For our application, we imagine that the parasite is persisting because of spatial structure and would go extinct under mass action – the large majority of hosts inhibit parasite reproduction because of the cargo. We address how selection acts on a weakly resistant mutation, a mutation that is not necessarily sufficient to provide positive parasite growth from inhibitory hosts alone.

It might seem valid to evaluate long term resistance evolution from a comparison of eigenvalues of wild-type and mutant growth, in which case the preceding figures could be used to infer evolution of alternative genotypes. However, we are considering resistance evolution in established populations at which density dependence is operating. When density dependence operates locally, as assumed here, it will have a different effect on parasite growth in cargo-free patches than in inhibitory patches. If the wild-type parasite cannot grow in inhibitory patches but the mutant can, the fecundities of both genotypes will be suppressed by density dependence in cargo-free patches but the mutant’s fecundity in the inhibitory environment will be unaffected (at least while rare). Eigenvalue calculations do not include these fecundity modifications. Our model of resistance evolution uses the system of equations in (3) but with the density-dependent carrying capacity enforced as in eqns (4) and (5).

This model was evaluated numerically for different combinations of fidelity (*α*), cargo-free patch size (*x*_0_), and genotype fecundities (*b*_*i j*_). We focused on the case of a wild-type (starting genotype) that was unable to grow in the inhibitory environment of cargo-bearing hosts (*b*_01_ = 0), and for which the mutant could grow in the inhibitory environment at some reduction in its ability to grow in the cargo-free environment (i.e., a trade-off was imposed between growth in the two patch types). The patterns across trials are qualitatively similar and easily comprehended (Fig. 3). Resistance could invariably evolve if it did not incur too much of a cost to growth in the cargo-free hosts.

**Figure 3.**
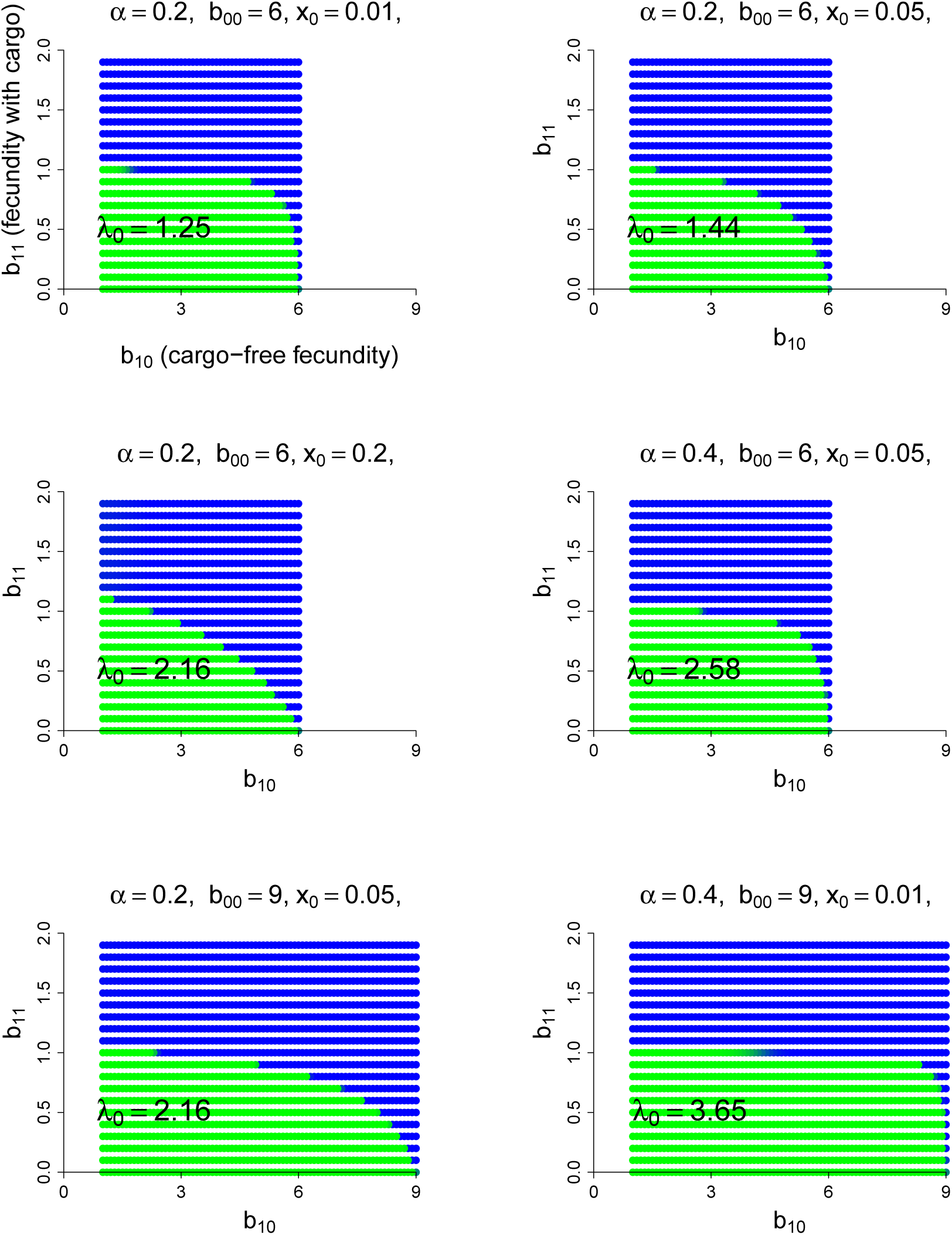
Parasite resistance to gene drive cargo, even if only partial, is favored if the cost to cargo-free fecundity is not too severe. Each panel represents evolution of wild-type versus mutant alleles under the set of parameter values given in the title. Patch-type fidelity is *α*, wild-type fecundity in cargo-free hosts is *b*_00_, and the patch size of cargo-free hosts is *x*_0_; wild-type fecundity in cargo-bearing hosts (*b*_01_) is zero in all panels. Each dot represents a different mutant allele whose fecundities in cargo-free hosts (*b*_10_) and in cargo-bearing hosts (*b*_11_) are given by its coordinates; axes in the upper left panel are labeled to assist recollecting the *b*_1*j*_. Green indicates that the wild-type was in a strong majority at the end of the trial, blue that the mutant was in a strong majority, and an intermediate color indicates that both alleles were moderately common. In the absence of competition from the mutant, the wild-type would persist for all conditions tested; its growth rate when rare is given as *λ*_0_ in the panel, from eqn (9). Trials were run for 1000 generations in which both alleles started equally abundant in both patch types. Carrying capacity was 10^6^ for all trials shown here.

Some ‘ecological’ patterns are evident. (1) A large effect of *α* – host patch fidelity – on evolution exists in some parameter ranges. Thus, to displace wild-type, the mutant is more sensitive to a reduction in cargo-free fecundity at larger *α* – until fecundity in cargo-bearing hosts is high enough that the mutant could sustain itself in those hosts alone (until *b*_11_ exceeded 1). This effect is evident when comparing lower portions of the top right and middle right panels in Fig. 3 and was seen in other trials (not shown). We interpret this effect as that higher values of *α* increasingly partition growth in the two patch types, so that any fitness loss in the permissive patch – the one sustaining the population – is increasingly penalized, but only to the point that the mutant parasite can maintain itself in cargo-bearing hosts.

(2) Patch size *x*_0_ also appears to have a large effect, but in the opposite direction as that of *α*: as *x*_0_ increases, resistance evolves more easily (i.e., tolerates larger reductions in cargo-free fecundity, *b*_00_ *-b*_10_). This effect is seen by comparing the first 3 panels of Fig. 3 which vary only *x*_0_. Although not shown, the trend does not continue at high values of *x*_0_, and the effect reverses. Indeed, mutants whose *b*_11_ > 1 are not favored at high *x*_0_ values if they suffer too much cost in *b*_10_. We do not pretend to grasp these patterns intuitively. One challenge in understanding these outcomes is that *x*_0_ has effects at different steps of the life cycle: with higher *x*_0_, the cargo-free patch has an increased carrying capacity and thus produces more progeny; some of these progeny return to patch 0 and the others go to the random pool. But the increase in *x*_0_ reduces the number of random-pool progeny that land in patch 1 and thereby reap the benefits of resistance.

## DISCUSSION

Genetic engineering has brought us to the brink of being able to introduce selfish genetic elements (known as gene drives) in countless species (Esvelt et al., 2014; Report, 2016; Collins, 2018). Although safety concerns and regulatory hurdles, and to some extent technical hurdles, have so far prevented the release of engineered gene drives, it seems inevitable that they will be released and eventually become a standard intervention for pest management and disease control—if they work as expected. Much excitement is about using gene drives for population suppression (Hamilton, 1967; Lyttle, 1977; Burt, 2003; Godfray et al., 2017; Lambert et al., 2018), but an alternative approach that may encounter fewer regulatory obstacles is to use ‘modification drives’ for spreading genetic cargo through a population (Sandler and Novitski, 1957; Gould, 2008; Gantz et al., 2015; Beaghton et al., 2017). In this latter application, the gene drive is a rapid and potentially harmless means of genetically engineering an entire population to carry a novel gene of interest. Thus, a mosquito that transmits *Plasmodium* might be targeted for eradication by a gene drive (Kyrou et al., 2018) or be targeted with a modification drive to spread a cargo that express anti-*Plasmodium* antibodies that block *Plasmodium* reproduction in the mosquito (Gantz et al., 2015).

Our study addressed the latter type of application and even assumed that the gene drive cargo had already spread in the host (e.g., mosquito) population. If this inhibitor fully suppresses the parasite in any host individual expressing the cargo—if it operates as expected—how might parasites persist despite our efforts to eradicate? Our emphasis here is on the possible contribution of spatial structure in the host population to parasite escape. Even when the gene drive successfully spreads the cargo to most of the host population, spatial structure combined with imperfect gene drive spread may leave pockets of cargo-free mosquitoes that allow the parasite to persist locally.

Our findings suggest that spatial structure in the host population can contribute to—indeed be sufficient to—enable parasite persistence against a cargo that would otherwise eradicate the parasite. Pockets of parasite persistence then foment the evolution of resistance to escape the cargo, unless resistance mutations cannot arise. The pockets need not be large, possibly representing a very small fraction of the range of the species targeted by the drive.

Our results follow work suggesting another reason that cargo-bearing gene drives may fail to eradicate parasites: the cargo frequency will begin to decline as soon as the gene drive carrier has reached its zenith in the host population (Beaghton et al., 2017). Thus, independent of spatial structure, any cost to carrying the gene drive element or cargo will select a loss of those elements once the gene drive spread has ended. An additional problem facing modification drives that use homing is that they target genomic sequences that are not essential to the host and thus have few selective constraints. Weakly selected sequences can tolerate variation that would block a gene drive. A clever solution to this problem is to use drives that target highly conserved sequences; to avoid harmful effects, they carry an insensitive cargo that replaces the target gene (Burt, 2003; Champer et al., 2019c). They must then carry two cargos, one for the modification, one to replace the target gene. Alternatively, use of multiple targets with CRISPR homing (multiple gRNAs) may limit resistance evolution (Champer et al., 2019d).

There are two requirements for parasite escape under the models studied here: spatial structure and genetic variation in cargo presence/expression that coincides with the spatial structure. Observation of a spatially structured population would indicate that the first requirement is met. Pre-existing genetic variation in the host population may even contribute to variation in cargo presence or expression, as when existing variation directly resists gene drive spread or affect cargo expression, and spatial structure of the variation would lead to both requirements being satisfied. But even if existing genetic variation does not affect gene drive evolution or cargo expression, any existing spatial structure may become a problem for genetic variation that evolves during gene drive spread. Gene drive spread can generate its own variation in cargo presence/expression by evolving as it follows demographic paths of reproduction in the host population (Beaghton et al., 2017). Such variation would be maintained by any spatial structure intrinsic to the host population.

The problem of parasite persistence and resistance evolution in response to a gene drive cargo has parallels with evolution of resistance in other contexts: antibiotic treatment of bacteria, use of chemical pesticides, and even genetically engineered ‘Bt’ crops. There is a widespread recognition that intermediate levels of pesticides and antibiotics favor the evolution of resistance (e.g., Gould and MacKenzie, 2002; Andersson and Hughes, 2012; Tabashnik and Gould, 2012; Neve et al., 2014; Gould et al., 2018). These problems are more often cast as stemming from temporal variation in dose rather than spatial variation, but the two types of heterogeneity are similar over the right time scales. In contrast, the practice of planting non-engineered strains among genetically engineered Bt crops to delay insect resistance evolution is explicitly one of destroying spatial structure (Tabashnik and Gould, 2012) and highlights the importance of spatial structure to the evolution of resistance. Gene drive cargo expression is presumably stable in time (for individual hosts and their dependants), at least for the short term, so parasite escape is primarily a problem of spatial variation rather than one of temporal variation.

The models leave many questions unanswered about the nature and magnitude of spatial structure required to enable parasite persistence. Indeed, it is the combination of spatial structure in conjunction with genetic variation that matters. The right kind of spatial structure might exist in one part of a species range, the appropriate genetic variation in another, yet the parasite be suppressed by the cargo throughout. Furthermore, the permanence of spatial structure in the host population will depend heavily on host dispersal patterns, and it is the combined movement over the the life cycle of the parasite that determines the relevant structure. For parasites that alternate between two host species (e.g., mosquito-borne diseases of humans), a highly structured mosquito species will not necessarily enable parasite persistence if the second host—humans—is sufficiently mobile on the right time scale. Issues such as longevity of the spatially structured patches, plus averages and variances of host dispersal distances may need to be explored in the context of specific applications before understanding the potential for parasite escape.

Evolution of resistance is perhaps the ultimate concern. If cargo can be engineered to be resistance-proof, spatial structure will be only a temporary setback as additional interventions are implemented. Spatial structure will have the largest impact in facilitating evolution of resistance to cargo when resistance can evolve only gradually, in small steps. In this case, parasite eradication might well be achieved were it not for structure, but the structure provides the nucleus for gradual evolution of resistance that ultimately enables the parasite to maintain itself on cargo-bearing hosts. Stacking multiple inhibitors in the same host individual (as proposed for malaria (Gantz et al., 2015)) may, in the ideal case, prevent stepwise resistance evolution. Here the concern is an evolutionary loss of part of the cargo, so that only single inhibitors operate in some hosts; spatial structure would then contribute to evolution of resistance.

It has been convenient to focus on cargo-free patches as the type of reservoir enabling parasite escape. An alternative—or additional—type of refuge to consider is patches of intermediate cargo expression, enabling the parasite to persist at some level and directly favoring resistance. Intermediate patches may occur with many levels of expression and may arise because of genetic background effects in the host species or may arise by mutation during the spread of the gene drive itself. Thus, if cargo expression is costly to the host, drives with reduced expression will spread even faster than drives with the original engineering. These mutant-drive cargoes will form their own spatial structure as they spread, which may then be maintained by intrinsic host structure. See (Weinstein et al., 2017) for an interesting study on the evolution and dynamics of spatial structure in competing bacterial strains. The engineering faces a delicate balance between cargo over-expression and cargo under-expression. Over-expression may impose a fitness cost that selects against the drive/cargo, whereas the under-expression risks facilitating parasite persistence and evolution of resistance. The effect of patch intermediacy on persistence may be evaluated for our 2-patch case in Fig. 1 merely by considering one of the two patches to be intermediate instead of extreme (e.g., cargo-free fecundity *b*_00_ would be depressed or cargo-bearing fecundity *b*_01_ would be greater than 0). In a sense, our two patch model describes a worst-case scenario for parasite persistence; we expect permissive conditions in the real world will be broader than our results suggest.

Subject to possible limitations of our analysis (see below), our findings can be tentatively used to inform implementation practices most likely to succeed. In any implementation, inhibitory cargo should be chosen so that resistance evolution is difficult (i.e., requires multiple steps) or impossible, as inhibitors that can be overcome with single mutations seemed assured of eventual failure. Anti-drive resistance evolution will also be a factor that should be considered and is likely to vary with design and even with host population characteristics (Champer et al., 2019a,b), but that concern is not any more important for spatial structure than without. Beyond that, there are a few design features that may facilitate parasite suppression and work to limit evolution of resistance.

- **Prevent emergence of spatially structured variation**. Existing spatially structured genetic variation in a wild species may be difficult to change, although inundating small areas with release of lab-reared strains may offer a temporary solution—as underlies the sterile insect technique (Klassen and Curtis, 2005; Dyck et al., 2005). However, spatial variation that arises from gene drive spread and evolution (Beaghton et al., 2017) may be reduced by gene drive release at multiple sites (North et al., 2013), especially sites of import, such as population centers: the different waves of advance will collide with other waves, reducing any spatial evolution from single release points. Releasing multiple, independent drives in the same population, as proposed to overcome resistance evolution within unstructured populations (Burt, 2003; Deredec et al., 2011), may limit the extent to which any area is completely free of cargo from at least one drive. As pointed out by a reviewer, killer-rescue systems may be far more susceptible to the generation of spatial structured variation than are homing drives; indeed, that is one of their oft-cited advantages.
- **Target areas of incomplete coverage**. Following gene drive spread, areas can be assessed for cargo presence and expression. Regions identified as having inadequate coverage can be targeted for additional interventions to offset the limited effect on parasites.
- **Consider gene knockouts as cargo**. Quantitative variation in cargo expression may be structured, just as with cargo-free hosts. Partial expression of cargo may be even more conducive to parasite resistance evolution than is a complete absence of cargo (by selecting intermediates). A gene drive system that knocks out a non-essential host gene required for parasite reproduction/transmission may be less subject to intermediate expression than is a cargo transgene and thus less likely to select resistance. A possible downside is that ablation of a non-essential host gene may carry larger fitness defects than does expression of a foreign transgene, and thus select resistance to the drive.

### Limitations of the models

The models analyzed here depicted spatial structure abstractly and used several other simplifications to achieve analytical tractability and comprehension: population regulation with a sharp threshold, deterministic dynamics, steady state analyses, and few host types. Parasite fecundity was abstracted to be as simple as possible. The models are best interpreted as augmenting intuition rather than formally capturing any natural process, as might be done with agent-based simulations (e.g. North et al., 2013, 2019). The results may thus be seen as to invite more formal analyses that include, at a minimum, explicit spatial structure but also small population sizes that would accrue near extinction. Despite these limitations, the results unambiguously point toward spatial structure as seriously impeding gene drive implementations using cargo – an otherwise promising use of gene drives. From our results, we conjecture that spatial structure can be sufficient to enable parasite escape from inhibitory genetic cargo in the host population, but we equally suggest that there are likely to be many details affecting the ease of escape and evolution of resistance. Indeed, it will be desirable to study models specific to the biology of an implementation before making critical decisions about engineering and sites of release (e.g., Eckhoff et al., 2017; Lambert et al., 2018; North et al., 2019).

## ACKNOWLEDGMENTS

We are very grateful to reviewers who offered their considerable insight to the biology of gene drives and helped correct some our misunderstandings. JJB and CHR acknowledge support from National Institutes of Health (R01GM 122079 and P20GM 104420). RG was supported by a WSU Honors College Distinguished Professorship and NSF Grant DEB 1354264. The content is solely the responsibility of the authors and does not necessarily represent the official views of the National Institutes of Health. The funders had no role in study design, data collection and analysis, decision to publish, or preparation of the manuscript.

